# Chromatic and Achromatic Contrast Sensitivity in the far Periphery

**DOI:** 10.1101/2025.03.22.644503

**Authors:** Norick R. Bowers, Karl R. Gegenfurtner, Alexander Goettker

## Abstract

The contrast sensitivity function (CSF) has been studied extensively since the 1960s, however most studies to date have focused on the central region of the visual field. The current study aims to address two gaps in previous measurements: First, it provides a detailed measurement of the CSF for achromatic and, importantly, also chromatic stimuli in the far periphery of up to 90 degrees of visual angle. Second, we describe visual sensitivity around the monocular/binocular boundary that is naturally present in the far periphery. In the first experiment, the CSF was measured in 3 different conditions: Stimuli were either Achromatic (L+M), Red-Green (L-M) or and Yellow-Violet (S-(L+M)) gabor patches. Overall, results followed the expected patterns established in the near periphery, but achromatic sensitivity in the far periphery was mostly under-estimated by current models of visual perception, the quick decay in sensitivity observed for red-green stimuli slows down in the periphery. The decay of sensitivity for yellow-violett stimuli roughly matches the decay for achromatic stimuli even up to the far periphery. For the second experiment, we compared binocular and monocular visual sensitivity at different locations in the visual field. We observed a consistent increase of visual sensitivity for binocular viewing in the central part of the visual field compared to monocular viewing, but this benefit already decreased within the binocular visual field in the periphery. Together, these data provide a detailed description of visual sensitivity in the far periphery. These measurements can help to improve current models of visual sensitivity and can be vital for applications in full-field visual displays in virtual and augmented reality.

## Introduction

The contrast sensitivity function (CSF) is a fundamental aspect of human vision, delineating the boundary between the visible and the invisible. Over the past decades, the CSF has been extensively studied and numerous factors have been identified that can influence contrast sensitivity (F. W. Campbell & Robson, 1968; Rijsdijk et al., 1980; N. V. S. Graham, 1989; S. M. Wuerger et al., 2002; Pelli & Bex, 2013; A. B. Watson & Ahumada, 2005; Carney et al., 2013). These factors include spatial and temporal frequencies (Robson, 1966; Kelly, 1979), variations in chromatic information (Mullen, 1985; S. M. Wuerger et al., 2002; Mullen & Kingdom, 2002; Hansen et al., 2009), stimulus size (Rovamo et al., 1993), stimulus orientation (Regan & Beverley, 1983; F. W. Campbell et al., 1966), and average luminance (S. Wuerger et al., 2020; R. Mantiuk et al., 2020) or the location of the stimulus in the visual field (Himmelberg et al., 2023).

Here, we extend these studies and investigate how the CSF changes in peripheral vision at large eccentricities. Due to the uneven distribution of photoreceptors (Österberg, 1935; Grünert & Martin, 2020) and processing resources in the retina (Strasburger et al., 2011; Stewart et al., 2020), visual acuity and processing varies significantly across the visual field. The entire human visual field encompasses more than 180 degree of visual angle (dva), a fact that is often overlooked (see Strasburger 2020 for a discussion of this common misconception). However, the far periphery of vision in particular has received less attention in research compared to the central region. To date, few studies have examined the CSF outside of the central region of the visual field, typically investigating eccentricities up to 45 dva (Mullen & Kingdom, 2002; Wright & Johnston, 1983; Baldwin et al., 2012; Virsu & Rovamo, 1979; Rovamo & Virsu, 1979; Chwesiuk & Mantiuk, 2019), or only measured selected stimuli (Hansen et al., 2009; Hess et al., 2008; Regan & Beverley, 1983). Moreover, most of these studies have focused on achromatic gratings, and few measure chromatic sensitivity in the periphery (Hansen et al., 2009; Mullen & Kingdom, 2002; Mullen et al., 2005). However, chromatic stimuli are of particular interest because it has been suggested that color vision becomes almost absent at eccentricities further than 40 dva (Ferree & Rand, 1919; Moreland & Cruz, 1959), especially for red-green (L-M cone) defined stimuli (Mullen, 1991; Mullen et al., 2005). In contrast, other studies have demonstrated that if the stimulus is large enough (Gordon & Abramov, 1977; Abramov et al., 1991; Hansen et al., 2009) or otherwise optimized for color perception (Noorlander et al., 1983), different hues can be perceived even up to 90 dva.

Based on these previous studies, there is a need for a systematic investigation of the CSF in the far periphery for both achromatic and chromatic stimuli. This need has become especially relevant as several models have been introduced to predict the CSF across the visual field in recent years (Bozorgian et al., 2022; R. K. Mantiuk et al., 2022; Ashraf et al., 2024; A. Watson, 2018). These models incorporate the factors discussed above in various ways. A limiting factor of these models is that they necessarily rely on fits to the available behavioral data, and then try to extrapolate into regions where data is not yet available. Due to the lack of comprehensive sensitivity measurements in the far periphery, a validation of these predictions has not yet been possible.

An important consideration for understanding visual sensitivity in the far periphery is the absence of binocular vision. Only the central ∼120 degrees of the visual field is binocular, while the remaining ∼80 degrees split between the left and right periphery is monocular (Howard & Rogers, 1996; C. Johnson et al., 2011). The difference between binocular and monocular contrast sensitivity has been investigated, but mostly in the center of the visual field by blocking one eye. Initial measurements of contrast sensitivity comparing monocular and binocular viewing showed enhanced visual sensitivity for binocular viewing by a factor of roughly [inine] (F. Campbell & Green, 1965; Blake & Fox, 1973). This would match a quadratic summation of the signal of both eyes (Legge, 1984). Although this approximation generally holds true, a recent meta-analysis has shown considerable variability in the estimated enhancement in sensitivity across studies. This scaling factor can be influenced by the spatial frequency, temporal frequency, and duration of the stimulus (Baker et al., 2018). However, whether this factor differs across the visual field has received only limited attention (C. Graham, 1931; Wood et al., 1992) and was not systemically investigated around the monocular/binocular boundary. The question remains: is there a consistent improvement up to the natural border between binocular and monocular vision in the periphery, or does the scaling factor depend on the eccentricity of the stimulus?

The current study has two objectives: First, we aim to acquire a detailed map of contrast sensitivity in the far periphery of vision extending up to 90 dva. We aim to accomplish this by providing comprehensive measurements of the CSF for both achromatic and chromatic stimuli across the visual field. Second, we seek to investigate how the difference in sensitivity between monocular and binocular vision varies across the visual field, with a particular focus on the natural boundary between binocular and monocular vision. Collectively, these measurements will provide critical insights into processing of peripheral vision and contrast sensitivity in the far periphery, while simultaneously providing useful measurements to refine and update current models of contrast sensitivity.

## Experiment 1: Contrast Sensitivity in the far periphery - Methods

### Human Subjects

Experiment 1 involved 30 naive psychophysics subjects, consisting of 3 males and 27 females, with an average age of 26 years (ranging from 19 to 35 years). All subjects were self-reported emmetropes or had their vision corrected through the use of contact lenses; glasses were not permitted to avoid occlusion in the periphery caused by the frames. Subjects self-reported no known visual or neurological disorders, and normal color vision was confirmed through the use of Ishihara plates (Ishihara, S. (2004). Ishihara’s tests for colour deficiency. Tokyo, Japan: Kanehara Trading Inc.). Experimental protocols adhered to the tenets of the Sixth Declaration of Helsinki and were approved by the local ethics committee of Justus-Liebig-Universität Gießen (LEK 2020 - 0015). Eye dominance was determined through the “hole-in-the-card” test, where subjects created an aperture with the hands held at arms length. (https://www.aao.org/eye-health/anatomy/eye-dominance). Most subjects (23/30) were right-eye dominant. Each subject completed 3 blocks of the experiment in a single session and returned for 3 separate 75-minute sessions. Subjects were randomly assigned to one of three experimental groups (Achromatic, Red-Green, Yellow-Violet), resulting in 10 subjects per condition.

### Experimental Setup

In order to project stimuli across the entire horizontal field, three Samsung Odyssey Neo monitors were set up side-by-side. These monitors are 1193×336mm in size, with a pixel resolution of 5120×1440. The refresh rate was set to 120Hz. The bit-depth of the monitors was configured to 10 bits per channel (bpc) by modifying the xorg.conf file on the computer to ensure fine-scale adjustments of contrast. These monitors support 10 bpc display without resorting to spatial or temporal dithering. Notably, these monitors have a 1-meter curvature, ensuring that each section of the screen is equidistant from the subject, who is positioned 1 meter away. With this setup, the monitors span roughly ∼208 dva of visual angle. The edges of the monitors are solid black, creating a 2.5 cm (1.4 dva) wide black strip that slightly disrupts the otherwise continuous display at ±34 dva from the center. Subjects’ heads were supported by a chinrest to minimize motion, and eye movements were recorded through the use of an Eyelink 1000 (SR Research Ltd., Ottawa, ON, Canada) to ensure proper fixation. The visual environment was relatively sparse in order to avoid any distractions during the experiment. A photo and schematic of this setup can be seen in Figure 9. Stimuli were displayed with a NVidia GE RTX 3070Ti graphics card. The operating system was Linux Ubuntu 22.04.3 LTS (”Jammy Jellyfish”). Stimuli were presented using MATLAB’s Psychtoolbox (Kleiner et al., 2007). MATLAB was run via the console with the options “-nosplash” and “-nodesktop” due to the IDE’s inability to display properly in 10 bpc mode.

**Figure 1:**
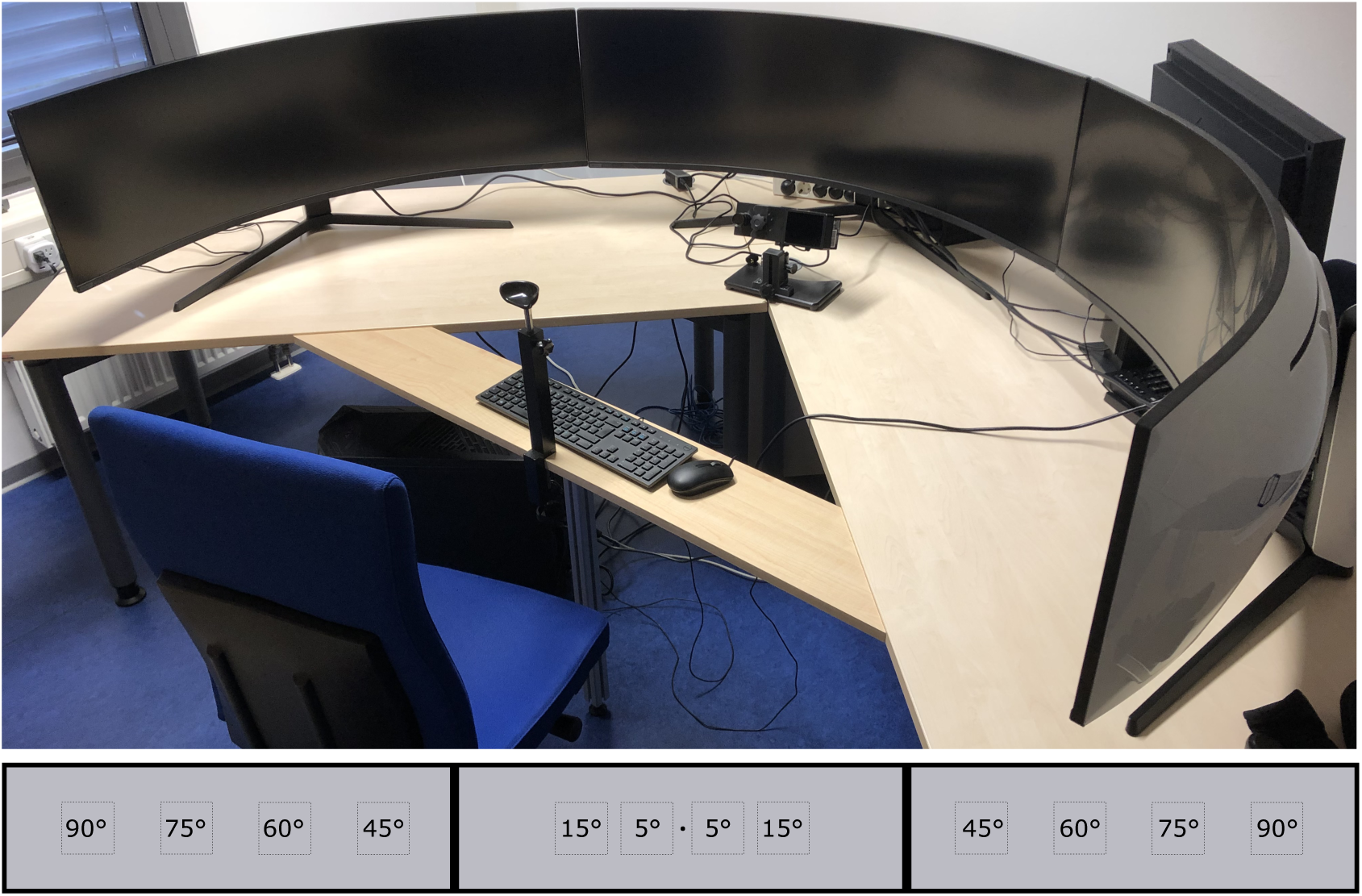
**Top**: A photo of the experimental setup. All three monitors together encompass approximately 208 dva. The borders of the monitors are relatively narrow (1.4 dva) and disrupt the otherwise continuous display at ± 34 dva from the center. Subjects were situated on the chinrest at a distance of 1 meter. An Eyelink 1000 (SR Research Ltd., Ottawa, ON, Canada) was used to ensure proper fixation. **Bottom**: Scale model of the monitor setup and stimuli locations for Experiment 1. The fixation dot is the small black dot in the center of the central monitor. The solid squares represent the 8 dva boundary of the Gabor patches (2 standard deviation) at each of the stimuli locations across the three screens.

#### Monitor Calibration

The monitors were calibrated with a Konica Minolta Spectroradiometer CS-2000A (Konica Minolta Holdings Inc., Marunouchi, Tokio, Japan). Each of the three monitors was measured and calibrated separately to compensate for small differences between the monitors. The primaries in CIE1931 xyY for the three monitors (left, center, and right) respectively were for the red channels, x: 0.678, 0.678, 0.678 (left, center and right monitor), y: 0.309, 0.308, 0.309, Y: 17.295, 17.099, 18.029, for the green channel, x: 0.274, 0.274, 0.274, y: 0.664, 0.662, 0.663, Y: 61.260, 61.390, 62.368, and for the blue channel, x: 0.158, 0.157, 0.158, y: 0.065, 0.065, 0.065, Y: 6.100, 6.128, 6.268. From these measurements, a transformation matrix (see Hansen & Gegenfurtner 2013) was computed to transfer the stimuli to a common DKL color space (Derrington et al., 1984). The Primaries in CIE1931 xyY with a white point at x: 0.322, 0.319, 0.319 (left, center, right monitor); y: 0.347, 0.345, 0.345, Y: 82.136, 81.462, 82.560.

### Experiment Design

#### Stimuli

For Experiment 1, CSFs were measured using a left/right 2 alternative forced-choice detection task. There were three different stimuli: Achromatic (L+M), Red-Green (L-M), and Yellow-Violet (S-(L+M)). Stimuli were displayed in DKL color space (Derrington et al., 1984). The Red-Green and Yellow-Violet stimuli were photometrically isoluminant, varying exclusively in hue while keeping luminance constant to match the background. Stimuli for Experiment 1 consisted of vertically oriented Gabor gratings that varied in spatial frequency (0.1, 0.3, 0.5, 1, 3, 5, 10 cpd) and location (± 5, 15, 45, 60, 75, or 90 dva). The phase of the Gabor was randomly chosen on each trial. The sine grating was 15 dva in size with an overlaid Gaussian filter that had a standard deviation of 4 dva, fading into the gray background. The large size of the Gabors was chosen to ensure at least a single cycle would be visible at each spatial frequency, as well as to maximize visibility in the far periphery. Stimuli were displayed with a Gaussian temporal profile; ramping up to maximum contrast, and then ramping down. The Gaussian profile had a standard deviation of 0.15sec with an exponent of 4. An example of the stimuli as well as the temporal ramp can be seen in figure 2. Stimuli contrasts were selected using QUEST+ in Matlab (A. B. Watson, 2017). Priors for the QUEST+ procedure were obtained through a pilot study with 3 trained psychophysics subjects.

**Figure 2:**
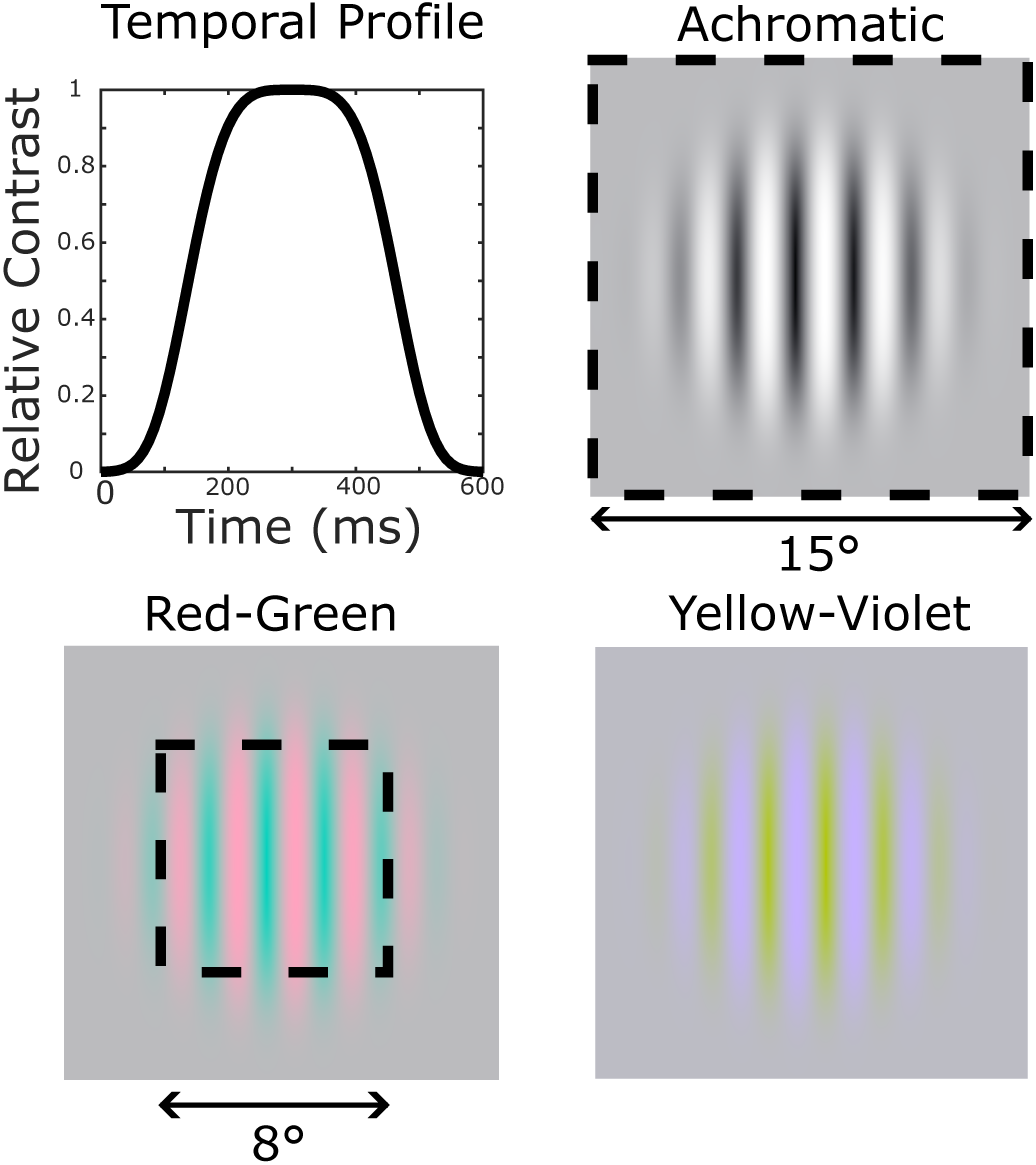
**Top Left**: The temporal profile of the stimuli. The stimuli would ramp on, reach max contrast, then ramp off over the course of 600ms. **Other panels**: example of the stimuli from the experiments (Achromatic, Red-Green, and Yellow-Violet) with the full 15 dva size of the texture as well as the 2 standard deviation 8 dva size highlighted by dashed lines on the Achromatic and Red-Green example, respectively. For this example the contrast of the gratings is set to 1.

#### Trial Procedure

Subjects were instructed to fixate on a small black dot (5 arcmin diameter) displayed on a gray background with a luminance of approximately 150 cd/m^2^. At the start of a trial the fixation dot would disappear, and the stimulus would subsequently appear on the display either to the left or the right of fixation, at one of the six possible peripheral locations. The stimuli would ramp on and off over the course of 600ms with the Gaussian profile described above. The fixation dot would then reappear, and subjects indicated which side of the display the stimulus appeared on using the arrow keys. A new trial would begin as soon as a response was given. Subjects were instructed to respond at their own rate, and were encouraged to rest and close their eyes briefly in between trials if they were experiencing fatigue. If subjects moved their eyes more than 2.5 degrees away from the central fixation dot location when the stimulus was on the screen, that trial was discarded and an identical trial appended to the end of the block.

Each block consisted of 10 valid repetitions of each combination of spatial frequency and eccentricity. Each subject completed three blocks of trials per session, with three separate sessions on different days. In total, subjects completed 90 valid trials for each combination of spatial frequency and eccentricity across all three sessions. Note that some combination of spatial frequency and eccentricity was not visible even at max contrast (e.g., 10cpd at 90 dva). These combinations were identified during the pilot experiment (where every combination of spatial frequency and eccentricity was shown) and excluded in Experiment 1. The exact combinations of spatial frequency and eccentricity that were included can be seen in the table below.

**Table 1:**
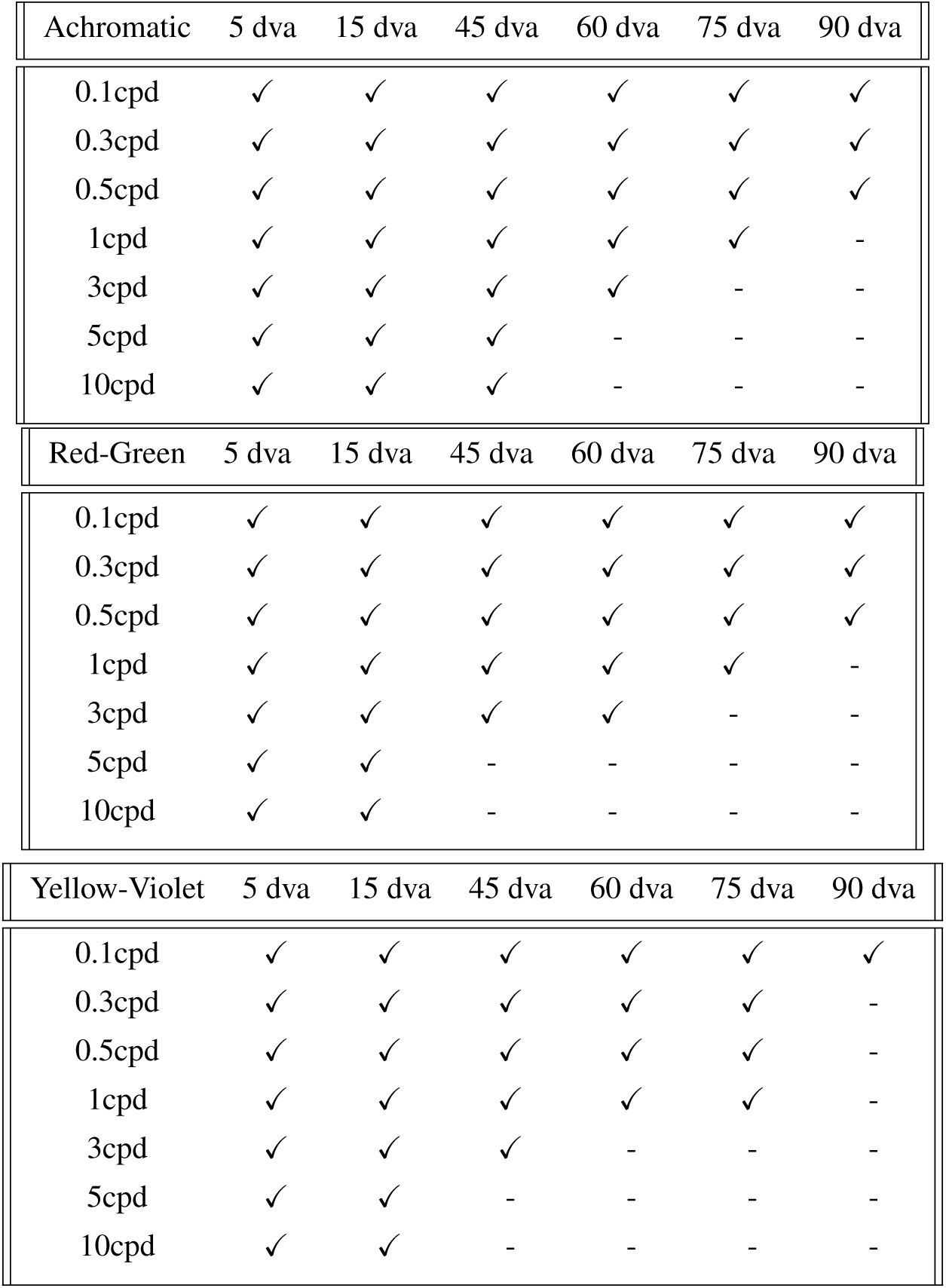
Table for each of the conditions (Achromatic, Red-Green, and Yellow-Violet) showing which combinations of spatial frequency and eccentricity were found to result in chance performance at max contrast during the pilot and removed from the experiment. Check marks indicate remaining combinations and dashes indicate those that were dropped. Combinations were included in Experiment 1 if even a single subject during the pilot exhibited performance above chance. Note that during Experiment 1, some of these combinations were found to result in chance performance for most subjects. We attribute this to the fact that the pilot was run with trained psychophysics subjects and Experiment 1 involved naive subjects. These “edge cases” that resulted in chance performance for Experiment 1 are discussed in more detail below.

### Data Analysis

To obtain contrast sensitivity values for each combination of spatial frequency and eccentricity we used a psychometric fit. The psignifit (Schütt et al., 2016) toolbox in Matlab was used to fit a Weibull function onto the 2AFC behavioral data. Contrast thresholds were defined at the 75% performance level. The presented contrasts were then transformed into cone contrasts by using the spectral distribution of the monitor primaries and the cone fundamentals from Smith and Pokorny 1975.

Once individual thresholds were acquired from the psychometric function, the data were fit with a contrast sensitivity function. For the achromatic condition, we expect the CSF curve to follow a band-pass shape and therefore used a log parabola to obtain a contrast sensitivity function using the following equation 1a (S. Wuerger et al., 2020):

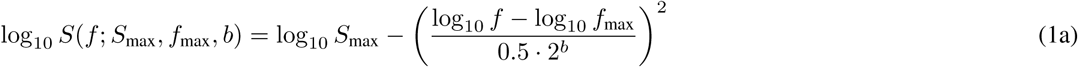

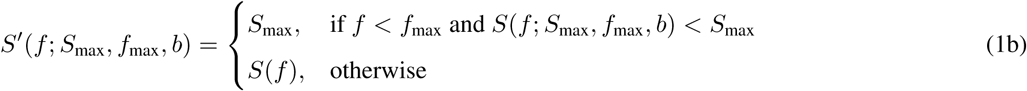

where *S_max_* is the peak sensitivity, *f_max_* is the peak frequency and *b* is the bandwidth of the curve. Since previous literature has found that the CSF for chromatic stimuli is generally low-pass in nature (Mullen & Kingdom, 2002; S. Wuerger et al., 2020), we used a truncated parabola with the same parameters, which was adjusted by equation 1b to encompass the low-pass nature of the CSF.

### Model comparison

To compare our newly measured behavioral data to current models of visual sensitivity, we used CasleCSF (v0.2.0) (Ashraf et al., 2024). The model is publicly available and we used parameters based on our stimuli and setup to create predictions. The parameters that were constant across all three experiments were: ‘t frequency’ = 0.3, ‘orientation’ = 0, ‘luminance’ = 150, ‘area’ = 64, ‘ge sigma’ = 4 and ‘xy background’ = [0.322, 0.347]. For the achromatic experiment we set the luminance modulation to 300 and ‘xy modulation’ was set to the same as ‘xy background’. For the red-greenish experiment, we set the luminance modulation to the same value as the background luminance and ‘xy modulation’ to [0.3970 0.3099]. For the yellow-violett experiment luminance modulation was again set to the background luminance and ‘xy modulation’ to [0.4042 0.5481]. With these inputs we then used our tested eccentricities as input for the ‘eccentricity’ parameter, and then used a a wide range of spatial frequencies to create the predicted CSF curves.

## Experiment 1: Contrast Sensitivity in the far periphery - Results

To estimate visual sensitivity in the far periphery, we used our setup to measure the contrast sensitivity function across the entire horizontal visual field up to 90 dva. There were three different types of stimuli: Achromatic, isoluminant Red-Green, and isoluminant Yellow-Violet.

### CSF Measurements

Raw individual contrast sensitivity measurements for each combination of spatial frequency and eccentricity are shown in figure 3. The figure presents three example observers in each condition (Achromatic, Red-Green and Yellow-Violett). It is important to note that the condition was a between-subject factor, so the data shown in the figure depicts nine unique observers and results cannot be readily compared between the achromatic and chromatic conditions. Several ‘edge cases’ occurred, particularly in the far periphery, where some subjects could detect the stimulus and some could not (e.g. compare observer RG9 and RG10 for 90 dva). Specifically, stimuli in both chromatic conditions at 90 dva were largely invisible, even at the lowest spatial frequency. Sensitivity measurements were obtainable for only 2/10 subjects in the Red-Green condition and a single subject in the Yellow-Violet condition for 90 dva eccentricity at the lowest spatial frequency, and these sensitivities were very close to the maximum contrast.

**Figure 3:**
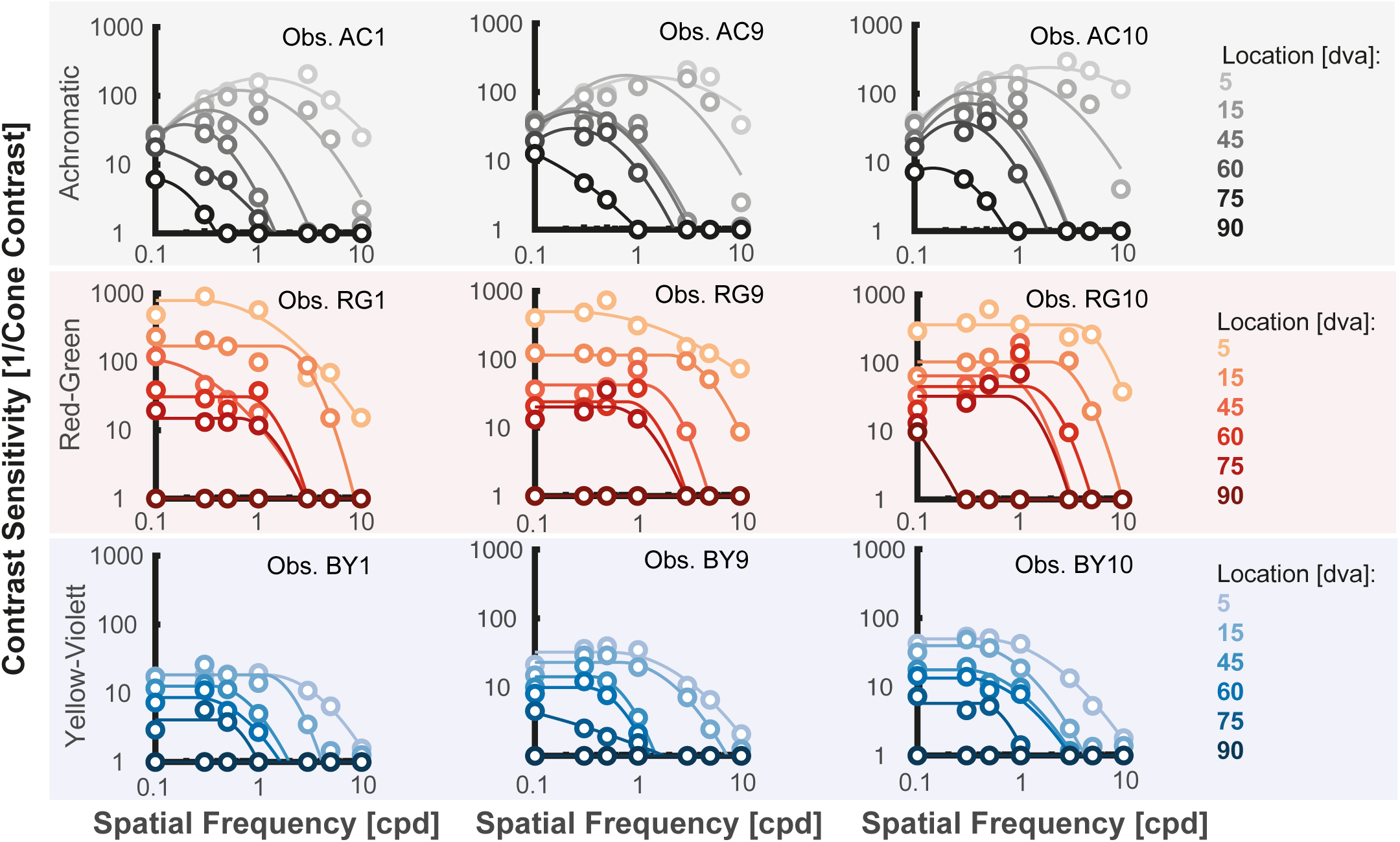
Individual contrast sensitivity measurements for 3 example observers are shown for each condition and across eccentricities. The top row shows data from the Achromatic condition, the middle row data from the Red-Green condition, and the bottom row data from the Yellow-Violet condition. Sensitivities from each of the conditions are shown in a different color: gray for Achromatic, red for the Red-Green condition, and blue for the Yellow-Violet condition. Darker shades indicate further eccentricities. Dots show the measured sensitivities and the lines show the fitted contrast sensitivity function. Please note here that some combinations for the far eccentricities were not measures, but we assumed no sensitivity due to pilot measurments (see Methods for more details). We used a log-parabola for the achromatic data, and a truncated log-parabola for the chromatic data

To better characterize the contrast sensitivity measurements, we fit a contrast sensitivity function to the individual data for each stimulus location for each subject (see lines in Figure 3). For the Achromatic condition, we used a log parabola fit to capture the expected band-pass nature of the data. For the two chromatic conditions, previous literature (Mullen, 1985; S. Wuerger et al., 2020) suggests we would expect a low-pass curve to better capture the data. To this end we fit a truncated version of the parabola on both the Red-Green and Yellow-Violet conditions.

To show the results across all of our 10 observers for each experiment (see Figure 4), we took the fitted curves for each individual and then averaged them per condition and location. Note here that the bends in the curves can happen due to individual variance in when stimuli become invisible. Note that while the figure shows the average of the CSF-fits, the curves would look nearly identical, if we would have first averaged the data and then fitted a new CSF curve. This is also true for the average parameter values of the CSF functions shown in the table below: the parameters obtained when averaging the sensitivity data first, and then fitting a single function to the averaged data, were always within one standard deviation of average of the individual parameter values.

**Figure 4:**
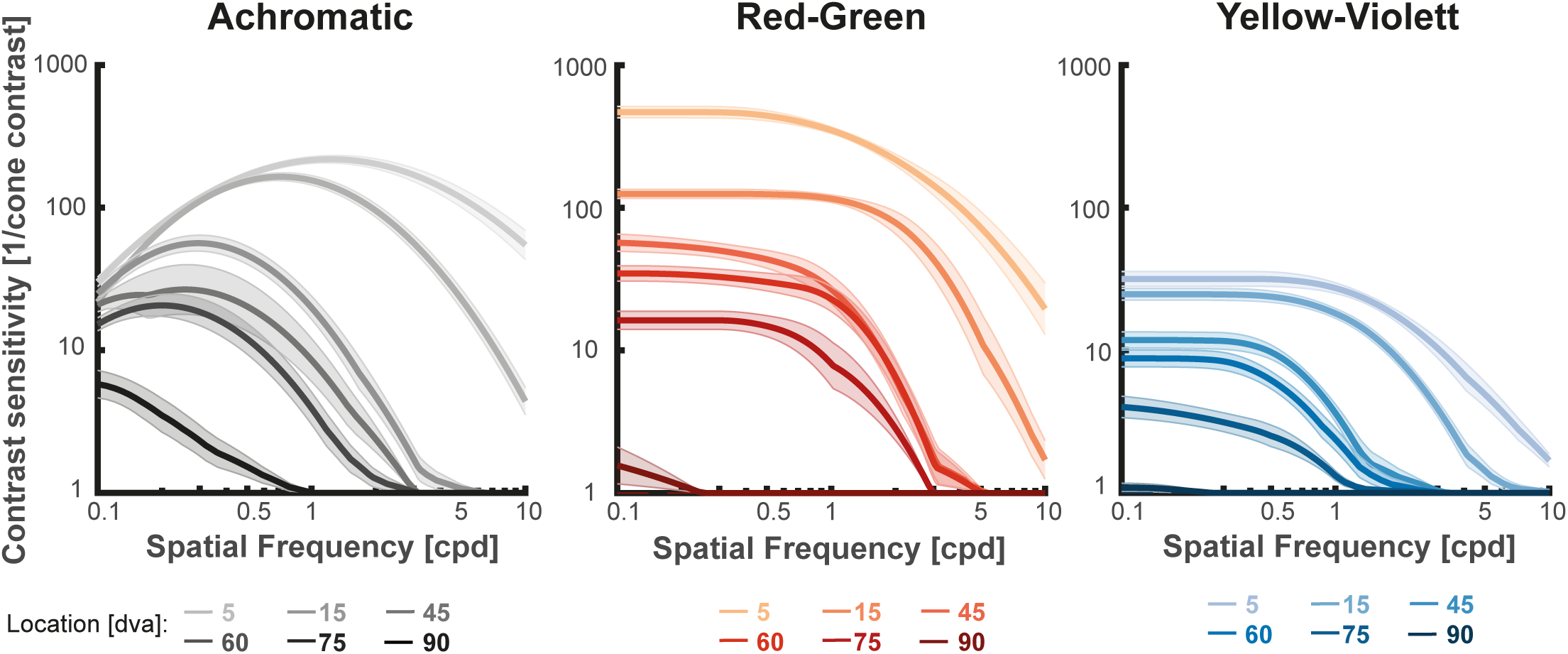
The average contrast sensitivity function across observers per condition and location. The solid line represents average of the respective individual fits and the shaded area shows the standard error between observers. The color scheme is the same as in figure 3.

The Achromatic CSF curves show a broadly consistent pattern: peak sensitivity decreases, and peak frequency shifts to lower spatial frequencies as eccentricity increases. The mean parameters of the fits are listed in Table 2.

**Table 2:**
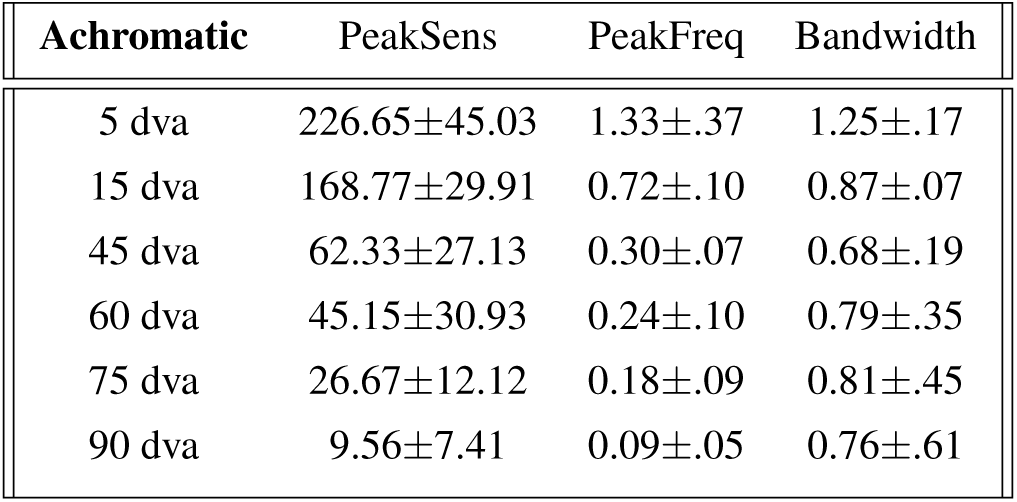
Table of parameters for the Achromatic condition. The measurements show the mean and standard deviation of the log-parabolic fit onto each individual subject’s data. The columns show the Peak Sensitivity, the Peak Frequency (the spatial frequency where peak sensitivity is obtained) and the Bandwidth of the curve.

| Achromatic | PeakSens | PeakFreq | Bandwidth |
| --- | --- | --- | --- |
| 5 dva | 226.65±45.03 | 1.33±.37 | 1.25±.17 |
| 15 dva | 168.77±29.91 | 0.72±.10 | 0.87±.07 |
| 45 dva | 62.33±27.13 | 0.30±.07 | 0.68±.19 |
| 60 dva | 45.15±30.93 | 0.24±.10 | 0.79±.35 |
| 75 dva | 26.67±12.12 | 0.18±.09 | 0.81±.45 |
| 90 dva | 9.56±7.41 | 0.09±.05 | 0.76±.61 |

A similar pattern to the Achromatic condition can also be observed in the Yellow-Violet condition, where the onset of the plateau is shifted toward lower spatial frequencies as eccentricity increases. In contrast, in the Red-Green condition the peak of the curve broadly remains similar across all eccentricities, even as sensitivity dropped with higher eccentricities. The parameters of the fits can be seen in table 3.

**Table 3:** Table of the parameters of the truncated parabola fit for each eccentricity for the two chromatic conditions. Measurements show the mean and standard deviation of the fit onto each individual subject’s data.

| Red-Green | PeakSens | PeakFreq | Bandwidth | Yellow-Violet | PeakSens | PeakFreq | Bandwidth |
| --- | --- | --- | --- | --- | --- | --- | --- |
| 5 dva | 486.82±144.60 | 0.64±.92 | 1.23±.60 | 5 dva | 33.58±11.68 | 0.64±.30 | 1.04±.31 |
| 15 dva | 131.43±28.75 | 1.37±.0.90 | 0.36±.37 | 15 dva | 25.37±7.71 | 0.63±.34 | 0.66±.52 |
| 45 dva | 62.08±24.15 | 0.60±.39 | 0.33±.53 | 45 dva | 12.57±4.32 | 0.33±.09 | 0.47±.39 |
| 60 dva | 36.88±12.63 | 0.70±.36 | 0.25±.49 | 60 dva | 9.40±3.13 | 0.27±.13 | 0.66±.48 |
| 75 dva | 17.79±7.76 | 0.60±.21 | 0.19±.20 | 75 dva | 5.70±3.78 | 0.22±.20 | 1.02±1.10 |
| 90 dva | - | - | - | 90 dva | - | - | - |

### Reduction in sensitivity across eccentricity

An interesting pattern in the data is that the loss of sensitivity is the largest for the Red-Green stimuli. To quantify this, we normalized the fitted peak sensitivities across the different locations by the peak sensitivity at the 5dva location (see Figure 5) and examined the normalized rate of sensitivity loss for each condition. When looking at the decay at 15 dva as a proxy for the initial drop in sensitivity as one moves away from the central visual field, the Red-Green condition dropped to around 29%±9%. This is much lower compared to the Achromatic condition (76%±14%) or the Yellow-Violett condition (79%±19%). This difference holds true across the visual field. When comparing the sensitivities for the 5 dva eccentricity with the 75 dva eccentricity, the Red-Green condition still shows the largest drop in sensitivity, with only 4%±2% of the initial sensitivity remaining. The Achromatic condition has the second largest effect, where 12%±5% of the initial eccentricity remains at 75 dva. The most stable sensitivity is present in the Yellow-Violet condition, where even at 75 dva, 18%±10% of the initial sensitivity still remains. When looking at the pattern of the decay across eccentricity, in the Red-Green condition there is initially a large drop in sensitivity which then slows down beyond 15 dva. For Achromatic and Yellow-Violet conditions, the loss in sensitivity is slower, but stays comparable across the whole horizontal visual field.

**Figure 5:**
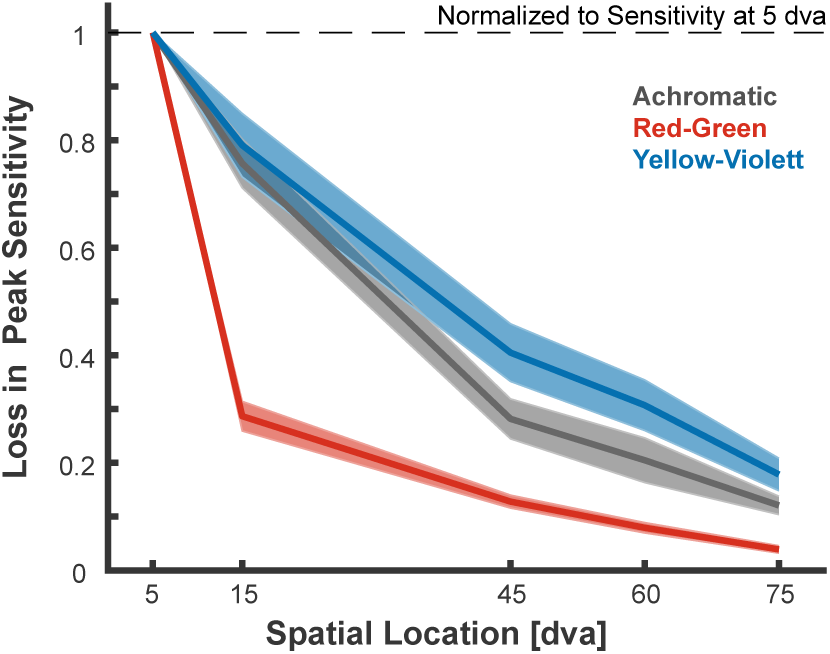
Shown is the average ratio between the peak sensitivity at the different eccentricities normalized to the peak sensitivity at 5 dva. The grayish line shows the reduction in sensitivity for the Achromatic condition, the red line for the Red-Green condition, the blue line for the Yellow-Violet condition. The shaded area depicts the standard error of the mean.

To illustrate how the rate of sensitivity loss was for each spatial frequency as a function of eccentricity, the data are replotted in Figure 6. There was a consistent decline in sensitivity with increasing eccentricity, however the decline becomes steeper at higher spatial frequencies. Figure 6 also highlights instances where some subjects were able to see the stimuli while others could not. This was particularly prevalent at the higher spatial frequencies as well as further eccentricities.

**Figure 6:**
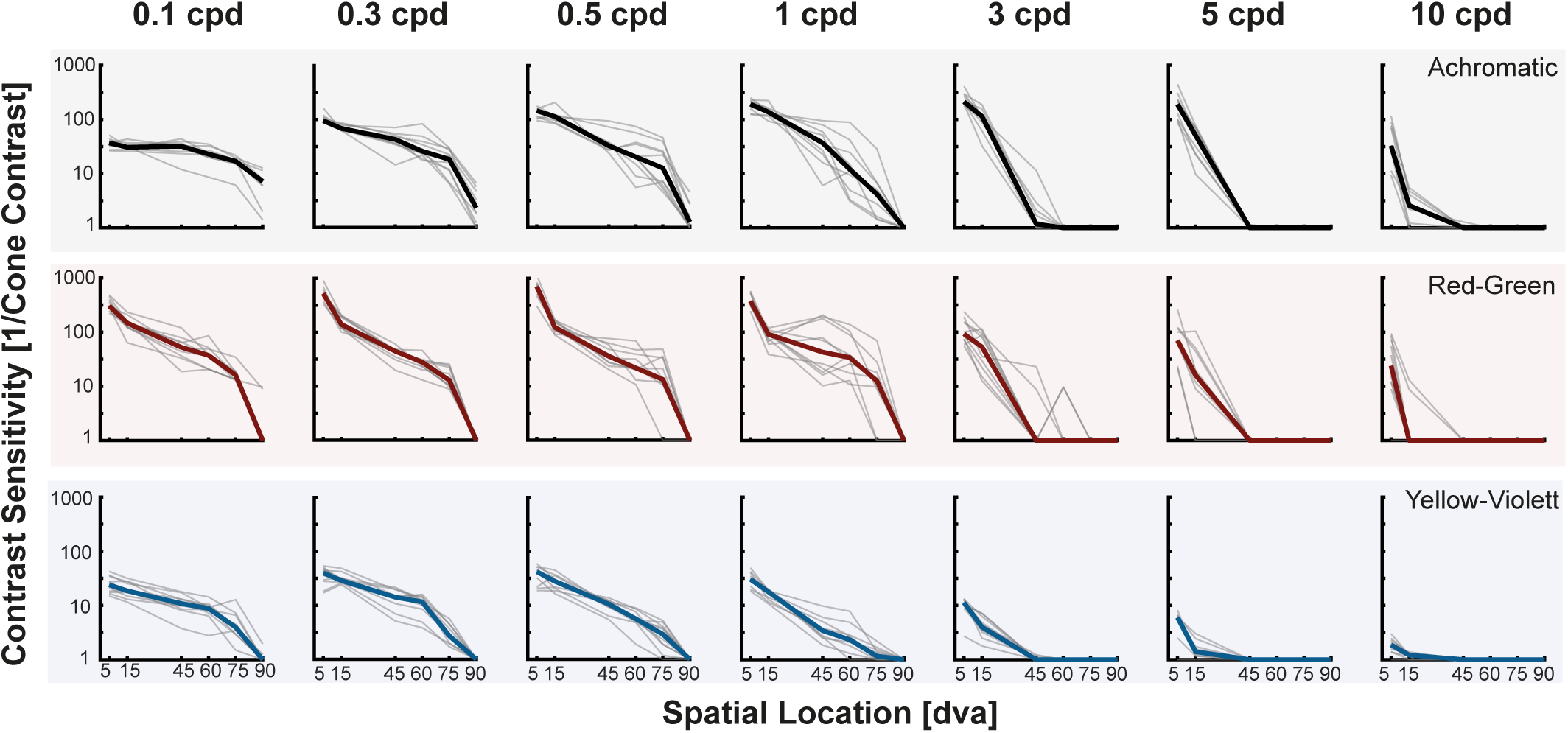
Individual (thin gray line) and median (thick line) data representing the loss of measured sensitivity with increasing eccentricity. Each row represents one of the experimental conditions (Achromatic, Red-Green, and Yellow-Violet; from the top-to-bottom respectively). Each column represents a different spatial frequency, with increasing frequencies from left-to-right.

To visualize the measured sensitivity in the periphery, we implemented a two-dimensional filter based on our sensitivity data (see A. Watson 2018 for a similar visualization). We used an image that had 30 pixels per degree and filtered it as it would span 180 x 60 dva. In contrast to previous approaches, we were now able to build a filter that correctly incorporates the decay of sensitivity for achromatic as well as chromatic information. We transform the original image into DKL color space, and then decompose each of the three axes (L+M, L-M, S-(L+M)) into different spatial frequency maps. The different spatial frequency maps were then weighted according to the predicted sensitivity for a given axes at a given eccentricity. The middle row of Figure 7, shows the original image decomposed into versions of the image reflecting the sensitivities at each of our measured distances. The bottom row shows continuous filtering of the image, where we estimate the sensitivity at each distance by interpolating between our measured CSF curves for each of the conditions.

**Figure 7:**
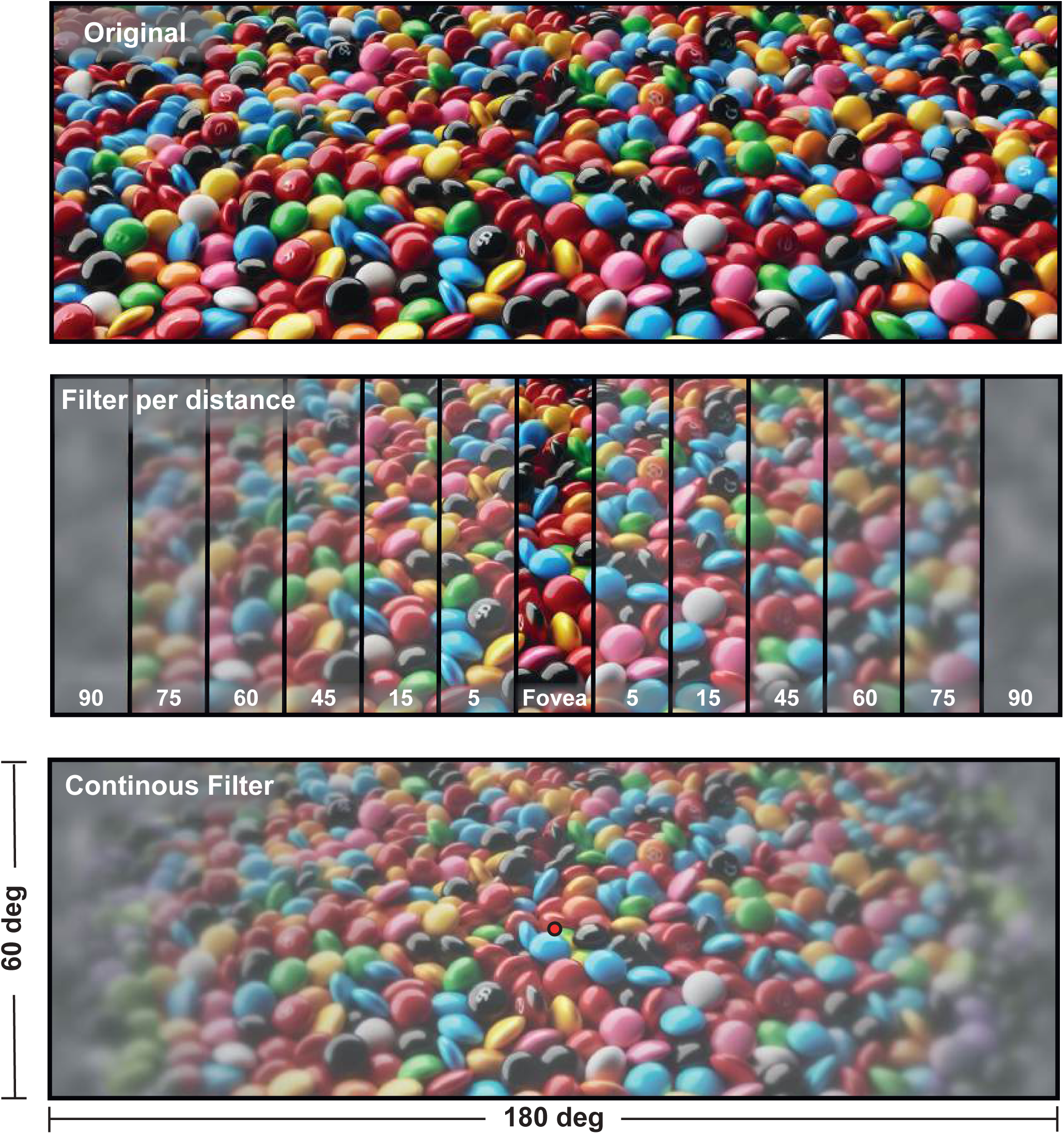
An image filtered according to our contrast sensitivity measurements. **Top** Original image created by an artificial intelligence. The image is setup in a way that it contains 30 pixel per degree and spans 180 x 60 dva. **Middle** The different sections of the image show how this part of the image would look like based on a filter matching our sensitivity measurements at the respective distance. For example, in this way it becomes visible that at 90 deg eccentricity low spatial frequency achromatic information is still visible, but color sensitivity is severely attenuated. Please note that the different sections reflect the measured distances which were not uniformly distributed. **Bottom** The same image now filtered continuously based on the respective distance to the central fixation point. For that, we extrapolated between our measured CSF curves to achieve a sensitivity estimate at each possible distance and spatial frequency for each of the three tested conditions.

### Comparison with Models

Several models have been proposed that predict contrast sensitivity in the periphery of vision (Bozorgian et al., 2022; R. K. Mantiuk et al., 2022; A. Watson, 2018). However, evaluating these models against psychophysical behavior in humans is challenging due to the relative lack of measurements in the far periphery. Rather, these models use the data from the more central region of the visual field and often extrapolate out to regions where no data exists. A recent model, castleCSF (Ashraf et al., 2024), is the only model that allows predictions for all of our tested conditions since it incorporates achromatic and chromatic sensitivity and eccentricity. CastleCSF is a data-driven, and physiologically inspired model that was fitted to a large data set of contrast sensitivity measurements of 18 studies providing measurements across varying factors. The model is publicly available and we used parameters matching our stimuli (see Methods for more details) to compare the current data from Experiment 1 to predictions from castleCSF. As with in original paper, we allowed one free additional parameter to linearly scale the castle CSF predictions for the 5 dva condition of the three experiments to match the overall levels of sensitivity measured in our paradigm close to the fovea and to then compare the behavior in the periphery. These predictions can be seen in figure 8 below. It is clearly visible that its predictive power largely diminishes at larger eccentricities. Especially for achromatic stimuli, the model predicts no clear change in sensitivity beyond 45 dva. In contrast, the decay in sensitivity for Yellow-Violett stimuli is underestimated.

**Figure 8:**
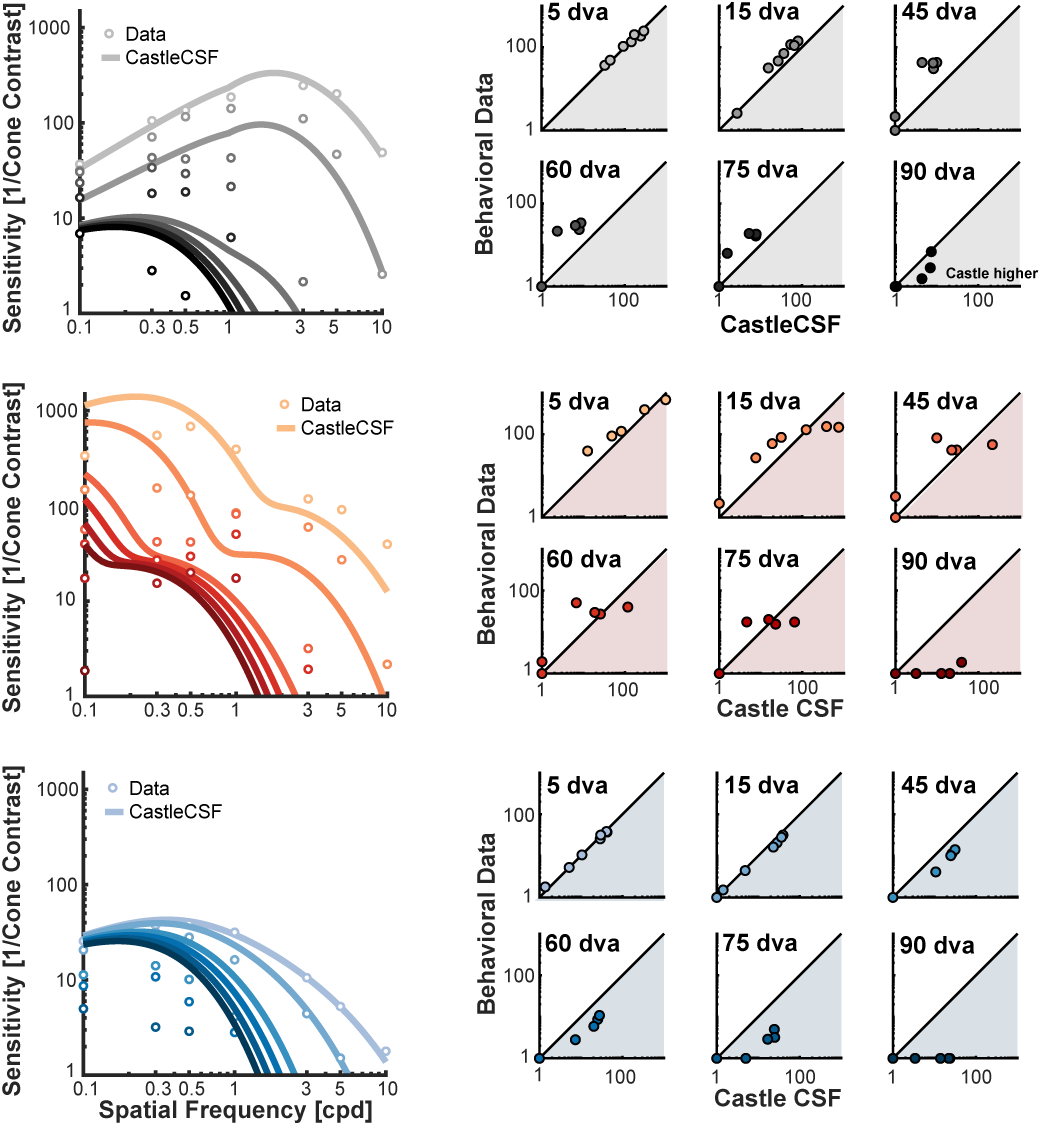
Comparison of current CSF measurements with predictions from CastleCSF (v0.2.0). **Left:** Contrast sensitivity measurements from the current study (dots) compared to predictions from CastleCSF (lines). The current data is the average sensitivity for each combination of spatial frequency and eccentricity across subjects. The predictions from CastleCSF are taken for all spatial frequencies between 0.1cpd and 10cpd for each eccentricity. **Right:** Differences between measured values and predicted values. The black horizontal line represents a perfect match between predicted and measured data.

## Experiment 2: Monocular vs. Binocular Sensitivity in the Periphery - Methods

### Human Subjects

For Experiment 2, a small subset of subjects from Experiment 1 returned to participate. In total, there were 10 subjects, consisting of 3 males and 7 females, with an average age of 26 years (ranging from 23 to 32). All participating subjects were right-eye dominant, as determined in Experiment 1. Experiment 2 consisted of two separate 75-minute sessions. In each session the subjects performed the task either monocularly, where the left eye was patched, or binocularly. The order in which subjects performed the two tasks was counterbalanced. Subjects had normal or corrected-to-normal vision and no known visual or neurological conditions. All subjects had normal color vision as determined in Experiment 1.

### Experiment Design

#### Stimuli

Stimuli for Experiment 2 were similar to those used in Experiment 1. However, to better assess the effects of monocular versus binocular viewing in the periphery, several notable changes were made. First, we only used achromatic stimuli. Second, subjects performed a ± 45 dva tilt discrimination task instead of a simple detection task to ensure that we could also measure sensitivity in the fovea as a baseline. Third, to reduce the number of trials, we constrained the spatial frequencies to 0.3cpd and 1cpd.

Stimuli were displayed at various eccentricities, some of which were relative to the binocular/monocular boundary measured for each subject, rather than being fixed. All stimuli were displayed at 5 different locations in the right visual field: Fovea, 20 dva, 10 dva left of the boundary, at the boundary itself, and 10 dva right of the boundary. Exactly how the boundary was measured is discussed in more detail below. The monitor setup was identical to that used in Experiment 1. The size of the stimuli, the Gaussian envelope, and the 600 ms ramp were kept the same as Experiment 1. Stimuli contrasts were selected using a Quest+ procedure (A. B. Watson, 2017), with priors for the QUEST+ procedure obtained through a pilot study involving a single trained psychophysics subject.

#### Trial Procedure

We used an adjustment procedure to establish the monocular/binocular boundary for each observer: before the first trial and after every 25th trial, subjects identified their monocular/binocular boundary by using the arrow keys to position a thin line (0.1 dva wide and 2 dva high) on the right side of the visual field at the boundary. Subjects were instructed to close their right eye and place the line at the boundary defined by the nasal orbit of the left eye and/or bridge of the nose as seen with the left eye. If subjects had difficulty closing one eye, they were allowed to cover their right eye with their fingers, provided that their fingers or nails were not visible to the left eye. Once the line was correctly located at the nasal boundary of the left eye, subjects pressed the space bar to continue.

In each trial, subjects judged the orientation of stimuli presented at one of the five locations on the right side of the visual field. During each trial, the procedure closely followed that of Experiment 1: the fixation dot disappeared, the target ramped up and down with the same Gaussian profile used in Experiment 1, and the fixation dot reappeared. However, instead of indicating which side of the screen the stimuli appeared on, subjects reported tilt using the arrow keys. Fixation was ensured in an identical manner to Experiment 1, using an Eyelink 1000 (SR Research Ltd., Ottawa, ON, Canada) to measure the gaze position while the stimuli was on the screen. If subjects moved their gaze more than 2.5 dva away from the fixation dot location while the stimuli was on the screen, that trial was discarded and appended to the end of the block to be repeated. The central fixation dot was shifted 5 degrees to the right to avoid potential overlap of the stimuli texture with the edge of the screen. In total, 120 trials for each combination of spatial frequency and location were collected, once in the binocular and once in the monocular condition.

**Figure 9:**
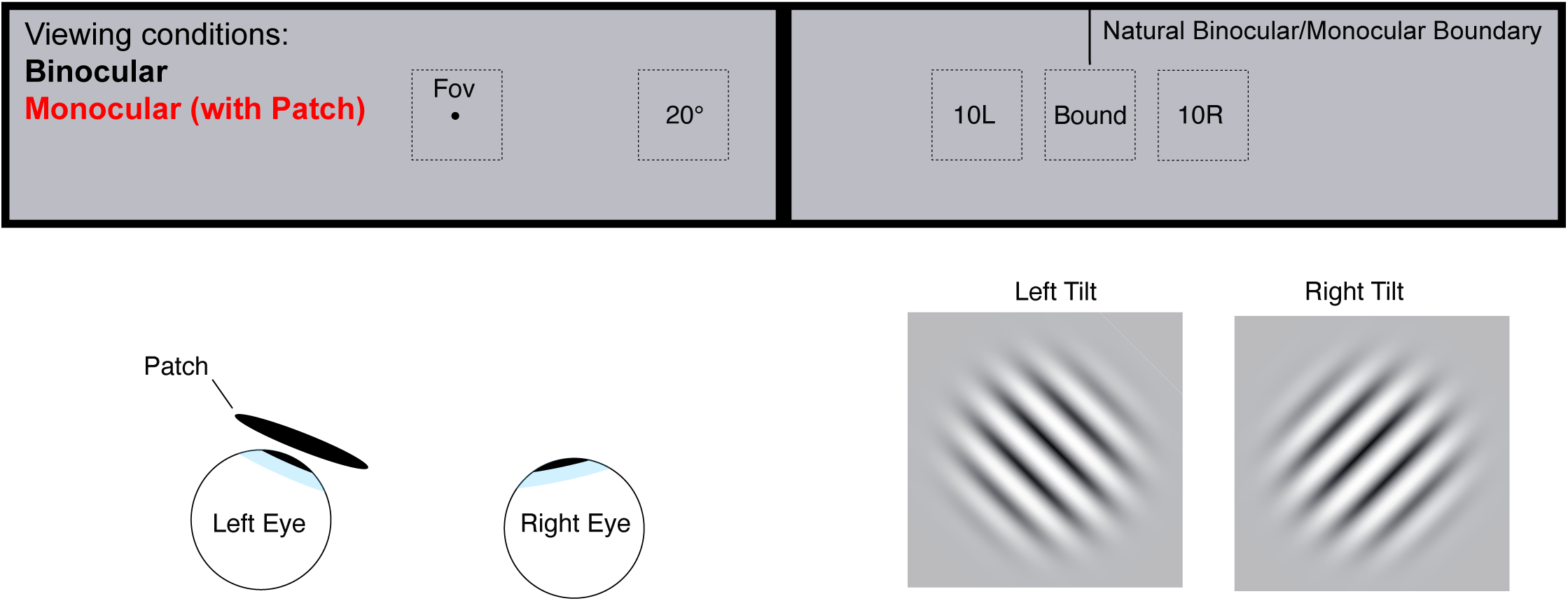
Scale model of the monitor setup and stimuli locations for Experiment 2. The fixation dot is the small black dot in the location marked Fov. This dot will disappear as the stimuli began ramping on. The fixation dot is shifted 5 dva to the right from the center of the monitor to avoid any possible overlap with the screen edge at location 10L. The solid squares represent the 8 dva boundary of the Gabor patches (2 standard deviation) at each of the stimuli locations across the screens. The left monitor (not shown) was blank and all stimuli were displayed to the right of fixation either monocularly (with an eyepatch over the left eye) or binocularly (without the eyepatch). The area marked Bound is the binocular-to-monocular boundary of the visual field identified per subject. Stimuli presented in this region would be half in the monocular visual field and half in the binocular visual field. The stimuli were ± 45 dva tilted Gabor patches and the subjects were instructed to report the orientation using the arrow keys.

### Data Analysis

In order to obtain contrast sensitivity measures from the raw data in Experiment 2, we used the same techniques from Experiment 1. We used the psignifit (Schütt et al., 2016) toolbox in Matlab to fit a Weibull function onto the raw behavioral data, with thresholds defined at the 75% performance level. These contrasts were transformed into cone contrasts in an identical fashion as Experiment 1. Since the goal of Experiment 2 was to compare performance between binocular and monocular viewing, it was not necessary to collect data on enough spatial frequencies to define an entire contrast sensitivity function. Therefore, we did not fit any function onto the data and relied on the raw threshold values for comparison.

To compare the monocular and binocular sensitivity, we computed the ratio between the binocular and monocular measurements. We then used a repeated measurement ANOVA to compare the ratios across two factors: spatial frequency (0.3 or 1 cpd) and the location of the stimulus (Fov, 20 dva, 10L, Bound and 10R). ANOVAs were performed in JASP and if Greenhouse-Geisser corrections were applied the corrected degrees of freedom are reported. Post-hoc tests were performed as well using the holm correction.

## Experiment 2: Monocular vs. Binocular Sensitivity in the Periphery - Results

In the second experiment, we wanted to systematically compare the visual sensitivity between monocular and binocular viewing conditions in the periphery. This comparison is especially critical in the far periphery, as there is a natural boundary in the visual field where the visual input switches between monocular and binocular signals. Here, we systematically investigated how the visual sensitivity in the temporal visual field changes when comparing binocular and monocular viewing. To test this, we introduced one condition in which the left eye was patched and so all stimuli were viewed monocularly and compared this to a condition where observers viewed the stimuli regularly with both eyes open.

We adapted the task used in Experiment 1 and turned it into a tilt discrimination task. To achieve an accurate measurement of the monocular and binocular conditions within the same observer, we concentrated our stimuli only on achromatic gratings with two different spatial frequencies: 0.3cpd or 1 cpd. Critically, stimuli were either presented in the fovea, or at one of 4 different locations to the right of fixation. Stimuli were either presented at 20 deg of eccentricity, or at one of three locations relative to the natural boundary between monocular and binocular visual fields: 10 deg to the left of this boundary (10L), on the boundary itself (Bound), or 10 deg to the right of this boundary (10R). The boundary between binocular and monocular vision is known to vary somewhat between individuals (Swanson et al., 2017; Glaser, 2017), so the exact location was set individually for each observer. Observers adjusted a thin line on the right side of the screen, so that it was barely visible when the left eye was blocked. The average location of the boundary was at 61±2.8 dva, and we repeated this procedure throughout the experiment to correct for potential head movements (see Methods for more details). With this approach and the size of our stimuli, the stimulus presented 10 deg to the left of the boundary is still completely in the binocular visual field. Furthermore, half of the stimulus for the Bound condition (i.e. at the boundary itself) should be in the binocular visual field.

We know from previous research that binocular vision leads to higher visual sensitivities than monocular vision. If this scaling factor is constant, we expect to see this improvement up to the 10L condition, then a potentially smaller improvement for the boundary condition (since only half the stimulus is still viewed binocularly), and no improvement for the 10R condition (which is only in the monocular visual field). Figure 10 shows the raw results for the two different spatial frequencies. In the Fovea and at 20 dva in the periphery, the binocular condition shows higher sensitivities than the monocular condition. However, for the 10L condition, which is still in the binocular visual field, this improvement seems to attenuate already.

**Figure 10:**
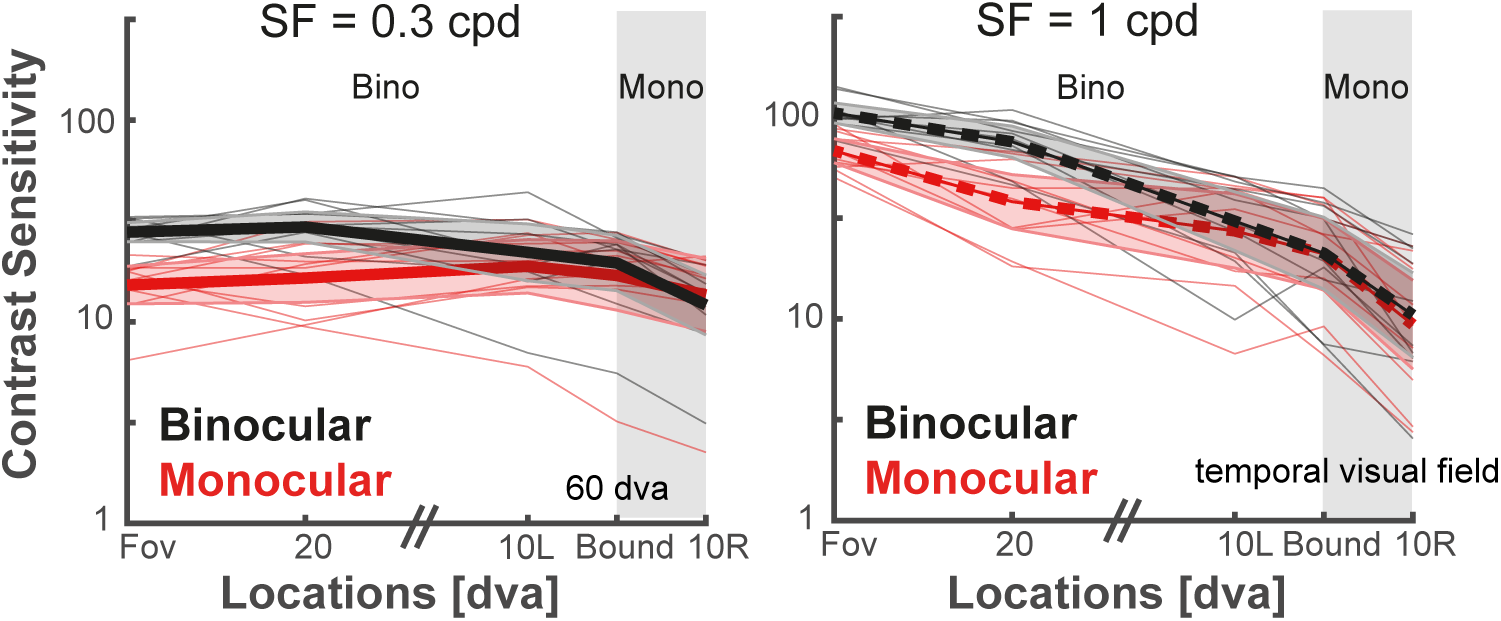
Comparison of binocular and monocular viewing. The panels show the contrast sensitivity for the stimuli with 0.3 cpd (left) and 1 cpd (right) spatial frequencies for the binocular (black) and monocular (red) conditions. The x-axis represents the different location of the stimuli. The stimulus was presented in the fovea or at 20 dva, or at various positions with respect to the monocular/binocular boundary: 10 dva to the left of the boundary, at the boundary, or 10 dva to the right of the boundary. The thick line shows the mean and the shaded area shows standard error. Thin lines represent individual subject data. The vertical gray shading indicates the locations that are outside of the binocular visual field.

To quantify the improvement, we computed the ratio between binocular and monocular sensitivity (see Figure 11). A ratio higher than 1 means higher sensitivity in the binocular condition, whereas a ratio of 1 would mean equal sensitivity in the two conditions. To assess whether the ratio is systematically affected by the location of the stimulus, we ran a repeated measurement ANOVA with the factors *spatial frequency* (0.3 or 1 cpd) and *location* (Fov, 20, 10L, Bound, or 10R). While the overall level of sensitivity was different between the spatial frequencies (see Figure 10A), for the relation between monocular and binocular vision there was no systematic influence of *spatial frequency* (F (1,9) =0.026, p = .875). However, there was a significant main effect of *location* (F(4,36) =13.808, p *<*.001) and no significant interaction (F(2.043,18.385) = 2.472, p = .111). Thus suggesting that there is a significant influence of the location on the scaling factor between monocular and binocular sensitivity that is consistent across the two spatial frequencies tested in our experiment. To investigate the influence of location in more detail, we used post-hoc t-tests and observed that the foveal and 20 dva condition were similar (p = .927), but they were significantly different from all other locations (all p values *<* .027 for foveal, and all p’s *<* .016 for the 20 dva location). This indicates that even for the 10L condition, which was fully in the binocular visual field, the binocular advantage was already diminished.

**Figure 11:**
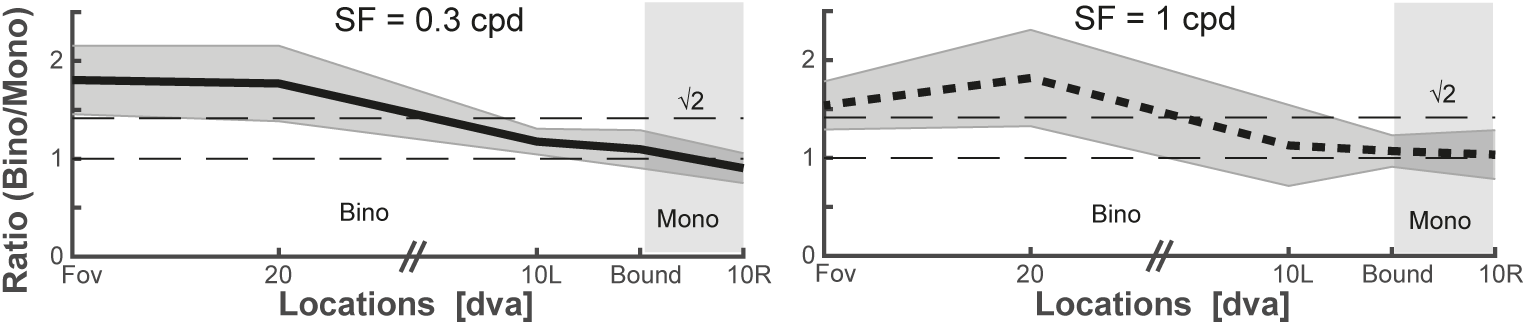
The panels shows the ratio between the binocular and monocular sensitivity across the different spatial locations for a stimulus with 0.3 cpd (left) and 1 cpd (right). The thick line shows the mean, the shaded area the 95 % CI. Dashed horizontal lines depict a ratio of 1, here there would be equal sensitivity between the two conditions, and 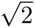 as a reference to previous literature.

To look into this further, we examined the direct comparison of monocular vs binocular sensitivities across our conditions. We observed clear differences for both spatial frequencies for the foveal and 20 deg peripheral condition (all t’s *>* 5.704, all p values *<* .001). For the 0.3 cpd condition, sensitivity was also significantly higher in the binocular condition for the 10L location (t(9) = 2.922, p = .017), but when correcting for multiple fall tests via Bonferroni (adjusted p-value: 0.005), this comparison did not survive the correction. No other comparison reached significance (all p values *>* .200). Thus suggesting that the scaling factor between monocular and binocular viewing diminishes in the periphery and is not constant across the visual field.

## Discussion

The current study provides a detailed description of visual sensitivity in the far periphery. In the first experiment, we measured the CSF across the horizontal visual field for achromatic and chromatic stimuli. As expected, we found that the CSF for achromatic stimuli shifted to lower peak frequency and lower peak sensitivity as a function of eccentricity. The pattern was similar for the Yellow-Violet condition, and the relative reduction in sensitivity was comparable to the Achromatic condition. For the Red-Green condition, the relative reduction in sensitivity was strongest and the shape of the CSF stayed roughly consistent. In general, stimuli with higher spatial frequencies become invisible, with only frequencies below 3cpd remaining visible beyond 45 dva. Comparison of our results with previous models that predict contrast sensitivity showed that visual sensitivity in the far periphery is typically overestimated for Achromatic and Red-Green conditions, and underestimated for Yellow-Violet conditions. In the second experiment, we compared sensitivity under monocular and binocular viewing with a specific focus on the natural border between monocular and binocular visual fields in the periphery. We observed that in the central visual field (up until at least 20 dva), there is a constant benefit for visual sensitivity for binocular over monocular viewing. However, this benefit seems to attenuate in the periphery, even within a part of the visual field that is still binocular.

### Comparison to previous sensitivity data

Qualitatively the pattern of results we find in achromatic and chromatic sensitivity across the visual field replicates previous findings. However, they go beyond previous results by providing empirical data on the sensitivity at peripheral locations between 45 and 90 dva.

For achromatic stimuli, we found that the CSF in the far periphery shifted to lower peak frequency and lower peak sensitivity as a function of eccentricity and this pattern continued up to 90 dva. This matches previous measurements made up to an eccentricity of around 45 dva (Pointer & Hess, 1989; A. Watson, 2018). Overall, our measured achromatic sensitivities in the central part of the visual field (up to 15 dva) were quantitatively larger than what has been observed in previous studies. However, we also used relatively large stimuli which where fixed in size in our experiments. Subjects unsurprisingly exhibit higher sensitivity to larger stimuli (Rovamo et al., 1993) and the loss of sensitivity in the periphery is understood to be largely caused by a loss of density in photoreceptors and retinal ganglion cells with increasing retinal eccentricity (A. B. Watson, 2014). It has also been known for many years that the brain devotes more resources to processing information in the central region of the visual field than the periphery (Rovamo & Virsu, 1979; Daniel & Whitteridge, 1961; Cowey, 1964), a phenomenon which can be captured by the Cortical Magnification Factor. The lower sensitivity in the periphery can be accounted for psychophysically by increasing the size of the stimuli according to this factor to make it match the sensitivity in the fovea (Vakrou et al., 2005; A. B. Watson, 1987). This is often done in such a way that the number of cycles visible in the Gabor remains constant. For the current study, we used a fixed size for our stimuli across all eccentricities and did not account for any type of magnification. This was done due to the excessively large size that would be required at the furthest eccentricities. Although this magnification factor is reliable and constant in the nearer periphery, it is unknown if this effect varies at very far eccentricities and our measurements show that with our relatively large stimuli, even in the far periphery at 90 dva, the stimuli with low spatial frequencies were still visible. In addition, and in contrast to many other studies, in experiment 1 participants also were viewing the stimuli binocularly, while most studies use monocular viewing. This is also known to increase sensitivity. See Baker et al. 2018 for a meta-analysis on this affect.

For the chromatic conditions, sensitivity in the central visual field was highest in the Red-Green condition, which aligns with expectations when comparing results in units of cone contrasts (Mullen & Kingdom, 2002; S. Wuerger et al., 2020). The Yellow-Violet condition exhibited the lowest sensitivity, which is likewise expected given the relative rarity of S-cones (Hunt & Peichl, 2014). The change in sensitivity as a function of eccentricity is also broadly consistent with previous results that compared Red-Green versus Yellow-Violet sensitivity from Mullen 2002 and Wuerger et al 1985 in the range of up to 15 dva where comparisons could be made. Similarly, we also observed that the loss of sensitivity was the strongest for the Red-Green condition, suggesting a stronger foveal specialization (see also (Mullen & Kingdom, 2002)). However, the loss in sensitivity became much more gradual in the far periphery (see Figure 5) and red-green stimuli with low spatial frequencies were still visible up to 75 dva. In contrast, the loss in sensitivity for the Yellow-Violet condition was much weaker and comparable to the Achromatic condition as described in (Mullen & Kingdom, 2002). This similarity holds up to 90 dva.

Our chromatic data showed that color vision is still present in the far periphery, at least to 75 dva. This is in contrast to previous reports that suggest color vision is strongly attenuated at around 40 dva (Ferree & Rand, 1919; Moreland & Cruz, 1959). A critical difference in the current study compared to these previous studies is that the stimulus was much larger in our data set. Earlier studies did observe cone-opponent color vision at more eccentric positions with larger stimuli (Abramov et al., 1991; Buck et al., 1998; Gordon & Abramov, 1977; M. A. Johnson, 1986; Hansen et al., 2009) and even up to 90 dva (Noorlander et al., 1983). Since our study also used comparably large stimuli, we confirmed the previous findings showing that with suitable conditions, color vision is possible even in the far periphery. While our results agree with some degree of functional specialization of color vision in the central part of the visual field, especially for L-M opponency (Mullen et al., 2005; Mullen, 1991), the ability to sense color in the far periphery must be due to a more general bias in the wiring of cone inputs to ganglion cells (Buzás et al., 2006).

### Comparison to visual sensitivity models

When directly comparing our data to recent models of visual sensitivity in the periphery (Ashraf et al., 2024; Bozorgian et al., 2022), we found interesting differences. For our results in the Achromatic condition, we compared our data to predictions from the modified Barten model from Bozorgian et al 2022. The results are not shown, but as for CastleCSF, this comparison was quite accurate for the nearer eccentricities, albeit the Modified Barten model generally predicted a more low-pass shape to the CSF curves in the near eccentricities. However, the modified Barten model predictions largely stayed consistent beyond 45 dva and predicted very little difference in the sensitivity at far eccentricities; whereas our data showed a more regular drop in sensitivity with a more band-pass shape and a shift of the peak frequency to lower eccentricities as eccentricity was increased.

The most complete comparison was possible for a very recent model of contrast sensitivity, CastleCSF (Ashraf et al., 2024). It attempts to account for many aspects of contrast sensitivity and was the only model that allowed us to make predictions for all of our tested stimuli. We found that sensitivity in the Achromatic and Red-Green conditions showed a much more gradual scaling and reduction in sensitivity in the periphery than what was predicted by CastleCSF. However, in the Yellow-Violet condition, CastleCSF predicted much higher sensitivity than our data showed, with little variation in the shape of the CSF curve as eccentricity increased. Thus, the current technique of fitting visual sensitivity models to the available behavioral data and then extrapolating the predictions to the far periphery cannot successfully capture the now available empirical data at larger eccentricities. This indicates that our measurements provide critical new behavioral data that can help to further improve such models.

### Limitations of the current measurements and next steps

There are two important questions related to the kind of sensitivity measurements we present here. First, one can test the limits of visual sensitivity for different types of visual stimuli presented on a display, and second one can try to differentially determine the sensitivity of the particular processing mechanisms in the visual system, in this case the putative achromatic and chromatic visual pathways. Our data are well suited to address the first question, but there are some caveats to keep in mind for the second question. There are significant differences between the light that reaches the eye and the way the light is shaped through its interaction with the optics of the eye. One crucial factor that can influence the measurements is chromatic aberration (Winter et al., 2016; Gawne & Banks, 2024). Chromatic aberrations are caused by the optics of the eye which cause rays of different wavelengths to take on different paths through the eye, which can degrade optical quality. In the case of isoluminant chromatic stimuli, chromatic aberration can also cause artifactual luminance contrast in the retinal image. There are two main types of chromatic aberrations. Longitudinal chromatic aberration (LCA) leads to different focal planes for lights of different wavelengths, see Gawne & Banks (2024)). This effect is roughly constant across the visual field (Jaeken et al., 2011; Rynders et al., 1998). Thus, it should not be able to explain the relative differences we see in visual sensitivity across different eccentricities for all of our stimuli. Second, there is transverse chromatic aberration (TCA) that is a misalignment of rays of different wavelengths on the retina, especially in off-axis locations (Winter et al., 2016). This effect is known to increase with eccentricity (Winter et al., 2016; Thibos, 1987; Flitcroft, 1989) and therefore could play a critical role explaining some of the effects we observe especially for our chromatic stimuli in the far periphery. While we cannot rule out that some of the sensitivity we measured for the chromatic gratings are due to chromatic aberration, there are three major reasons why we believe our measurements are close to reflecting the actual sensitivity of the visual processing mechanisms and not just optical aberrations. First, the effect of TCA should be in particularly strong for stimuli with higher spatial frequencies, since the abberations can produce stronger overlap between the different projections. However, similar to previous measurements (S. Wuerger et al., 2020), our chromatic data show a low-pass behavior of the curve. The highest sensitivity is usually at the lowest spatial frequencies, where the effect of TCA is weakest. Second, while we still see reliable visual sensitivity for low spatial frequencies for achromatic stimuli at 90 dva, only very few subjects had significant chromatic sensitivity at this eccentricity. Since the strength of TCA seems to be linearly related to eccentricity (Winter et al., 2016; Thibos, 1987), the effect of TCA should be even stronger at further eccentricities, which then cannot explain why there the sensitivity for our chromatic stimuli is suddenly reduced. Third, the level of peak sensitivity in cone contrast is highly comparable for achromatic and red-green isoluminant stimuli, for example at at 75 dva (see Figure 4,. Thus, it seems unlikely that the sensitivity in the Red-Green condition can be fully explained by a small amount of additional luminance information through chromatic aberration, as the sensitivity is comparable to a stimulus actively modulated in luminance. Together this suggests that while we mostly targeted to measure the visual sensitivity for stimuli presented on a display, our data should also provide a quite good estimate of the actual underlying sensitivity mechanisms when taking into account optical aberrations. This should be in particular true for the red-green stimuli (so the L-M mechanisms), as TCA here should be also weaker due to the closer similarity in terms of the wavelengths of the stimulus compared to the yellow-violet stimulus.

The current data set gives a relatively coarse measurement of contrast sensitivity in the far periphery. For example, although the loss of sensitivity with increasing eccentricity follows a relatively linear trend (for the Achromatic and Yellow-Violet conditions, less so for Red-Green condition), it is not perfectly linear. It is possible that more fine-scale measurements in the intermediary or far periphery would show this trend to reliably deviate from linear. Furthermore, obtaining reliable contrast sensitivity measures is a rather slow and laborious process using conventional psychophysics. We decided to obtain our measurements using an already established, tried-and-true method. However, newer methods of obtaining contrast sensitivity with less trials are available that rely on established assumptions on the shape of the CSF (Rosén et al., 2014), which based on our data are now available for the far periphery. These newer methods could be used to dramatically speed up data collection and allow for much finer scale measurements across the visual field. A more detailed measurement also seems critical since our data so far only capture the horizontal visual field. Our data might have limited usefulness in predicting contrast sensitivity along the vertical dimension. Previous research has consistently found a horizontal–vertical anisotropy, that is sensitivity along the vertical dimension was found to be worse than along the horizontal meridian(Rovamo & Virsu, 1979). Furthermore, there is also a vertical asymmetry, with worse sensitivity in the upper hemifield compared to the lower (Edgar & Smith, 1990; Previc, 1990; Hanning et al., 2022). Interestingly, this performance seems to worsen as the eccentricity is increased (Carrasco et al., 2001). It is unknown to what extent these anisotropies may persist in the far periphery as most research to date has stuck with relatively small eccentricities.

### Monocular vs. binocular vision across the visual field

Most previous studies measuring contrast sensitivity are done under monocular viewing conditions. However, for the central 120 dva of our visual field, naturally we have binocular input. Visual sensitivity is known to be higher under binocular viewing conditions compared to monocular, and in the fovea, this factor was classically thought to be around 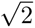 (F. Campbell & Green, 1965; Blake & Fox, 1973), which would match a quadratic summation of the signal of both eyes (Legge, 1984). However, more recent data show that this factor seems to be a slight underestimation and that it also differs due to some characteristics of the stimulus. In a recent meta-analysis, Baker et al 2018, showed that the scaling factor seems to be slightly higher for lower spatial frequencies, and can also be influenced by the temporal frequency or the duration of the stimulus. This fits well with our observations, as our measured scaling factor in the fovea was slightly higher than 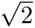 (see Figure 11, but we also used stimuli with low spatial frequencies.

In addition, our data show that not only the characteristics of the stimulus, but also its location in the visual field can affect the difference between binocular and monocular viewing. We observed that even within the binocular visual field, at eccentricities of about 50 dva, the difference in sensitivity between monocular and binocular viewing is attenuated. Our results are in line with previous work showing that the benefit of binocular over monocular viewing is dependent on the spatial location (Wood et al., 1992). It has also been suggested that the benefit is the largest in the central part of the visual field, where the two viewing fields overlap the most (Esterman, 1982). Our data seem to suggest that when predicting visual sensitivity based on monocular viewing data, it is fair to assume a roughly constant scaling up to 20 dva eccentricity and then an attenuation of the benefit towards the binocular/monocular boundary.

### The importance of understanding peripheral sensitivity

The high levels of sensitivity we still observe even at far eccentricities highlights the importance of peripheral vision. Peripheral vision plays a vital role in everyday tasks such as driving (Owsley & McGwin, 2010; Huisingh et al., 2014) and walking (McManus et al., 2017) and is essential for directing attention and foveation to stimuli of interest around the environment (for a comprehensive review on the use of peripheral vision in real-world situations, see Vater et al 2022). Understanding contrast sensitivity in the far periphery has significant implications for emerging technologies, such as augmented and virtual reality as larger fields of view become the standard. In particular, a thorough measurement of peripheral contrast sensitivity would benefit the development of foveated rendering algorithms (Meng et al., 2018; Guenter et al., 2012). These algorithms require reliable predictions of contrast sensitivity in the peripheral visual field from models to properly render the visual environment, and the current data show that these models do not accurately account for human psychophysical results in the far periphery. However, these models could be updated using the current data set to give more accurate predictions.

## Conclusion

Although visual sensitivity changes drastically with eccentricity, so far only limited behavioral data were available to describe the exact changes, especially in the far periphery. Our data now provide a detailed description of achromatic and chromatic visual sensitivity for eccentricities up to 90 dva. Qualitatively, we found that the pattern of changes in sensitivity seen at small and intermediate eccentricities seem to continue into the far periphery. However, there are a number of interesting differences: (1) Achromatic sensitivity in the far periphery is underestimated by current models of visual perception. (2) The quick decay in sensitivity for red-green stimuli slows down in the far periphery, (3) the decay for yellow-violet is comparable to achromatic stimuli even in the far periphery, and (4) the difference in sensitivity between binocular and monocular vision is not consistent across the visual field and is already attenuated within the binocular visual field at roughly 50 dva. Together these measurements provide a large step towards a better description of visual sensitivity in the far periphery, which is critical for reliable models of the visual system and for using visual models for full-field virtual or augmented reality displays.

## Acknowledgments

The authors want to thank Nike Leucker for help with data collection. NRB was supported by a Humboldt-Fellowship. KRG and AG were supported by the project numbers SFB/TRR 135 Project A1 (Project Number: 222641018). This project was supported by Meta Platforms, Inc.

## Data availability

The raw sensitivity data is available at OSF: https://osf.io/d3hw9/

